# Demonstration of *in vivo* engineered tandem duplications of varying sizes using CRISPR and recombinases in *Drosophila melanogaster*

**DOI:** 10.1101/2023.01.08.523181

**Authors:** David W. Loehlin, Georgia L. McClain, Manting Xu, Ria Kedia, Elise Root

**Affiliations:** Biology Department, Williams College, Williamstown, MA 01267

**Keywords:** Tandem duplication, copy-number variation, segmental duplication, recombinase, CRISPR, Alcohol dehydrogenase, *Adh*, genome engineering

## Abstract

Tandem gene duplicates are important parts of eukaryotic genome structure, yet the phenotypic effects of new tandem duplications are not well-understood, in part owing to a lack of techniques to build and modify them. We introduce a method, Recombinase-Mediated Tandem Duplication (RMTD), to engineer specific tandem duplications *in vivo* using CRISPR and recombinases. We describe construction of four different tandem duplications of the *Alcohol Dehydrogenase* (*Adh*) gene in *Drosophila melanogaster*, with duplicated block sizes ranging from 4.2 kb to 20.7 kb. Flies with the *Adh* duplications show elevated ADH enzyme activity over unduplicated single copies. This approach to engineering duplications is combinatoric, opening the door to systematic study of the relationship between the structure of tandem duplications and their effects on expression.

## Introduction

Tandem duplicate genes are a prevalent feature of genomes, with at least 17% of *Drosophila melanogaster* genes occurring in tandem clusters (Ashburner *et al*. 1999). Duplication of an entire gene produces a redundant copy, but it may also alter the phenotype through changes in gene expression. Understanding the expression outcome of tandem duplication mutations could be useful for understanding the evolutionary trajectory of duplicated genes and for rational design of gene expression in genetic engineering (Lan and Pritchard 2016; Birchler and Yang 2022; Loehlin *et al*. 2022). Although a simple prediction holds that duplicating a gene will double the gene expression level, current studies suggest that deviations from this two-fold hypothesis are frequent for transgenic and naturally occurring tandem duplicates (Cardoso-Moreira *et al*. 2016; Lan and Pritchard 2016; Loehlin and Carroll 2016; Hayward *et al*. 2017; Rogers *et al*. 2017; Konrad *et al*. 2018; Loehlin *et al*. 2022). Further evidence that tandem duplicated genes may not express independently from one another comes from the observation that tandem genes are co-regulated in important developmental processes (Levo *et al*. 2022). Understanding when, how, and why such deviations occur will be critical for developing a theory of tandem duplicate gene expression. To systematically investigate these questions, flexible techniques for creation and modification of tandem duplicate genes will be required.

In the wild, tandem duplication mutations are thought to originate by ectopic homologous recombination, in which a crossover or repair event occurs between non-allelic but otherwise identical sequences (Carvalho and Lupski 2016). Emulating ectopic crossovers has promise for engineering tandem duplications. However, the key first step, where a double-strand-break occurs in one chromosome homolog and not the other, is not easily achieved with current endonuclease-based technologies such as CRISPR-Cas9. We speculated that a two-step approach could work (Figure 1): In the first step, two modified chromosome homologs are generated, by separately inserting marked sequences to the left, and to the right, of the segment to be duplicated. The two asymmetrically modified homologs would then be introduced to the same cell by genetic crosses, followed by induction of ectopic crossing-over between the modified sites using a sequence-specific endonuclease.

**Figure 1.**
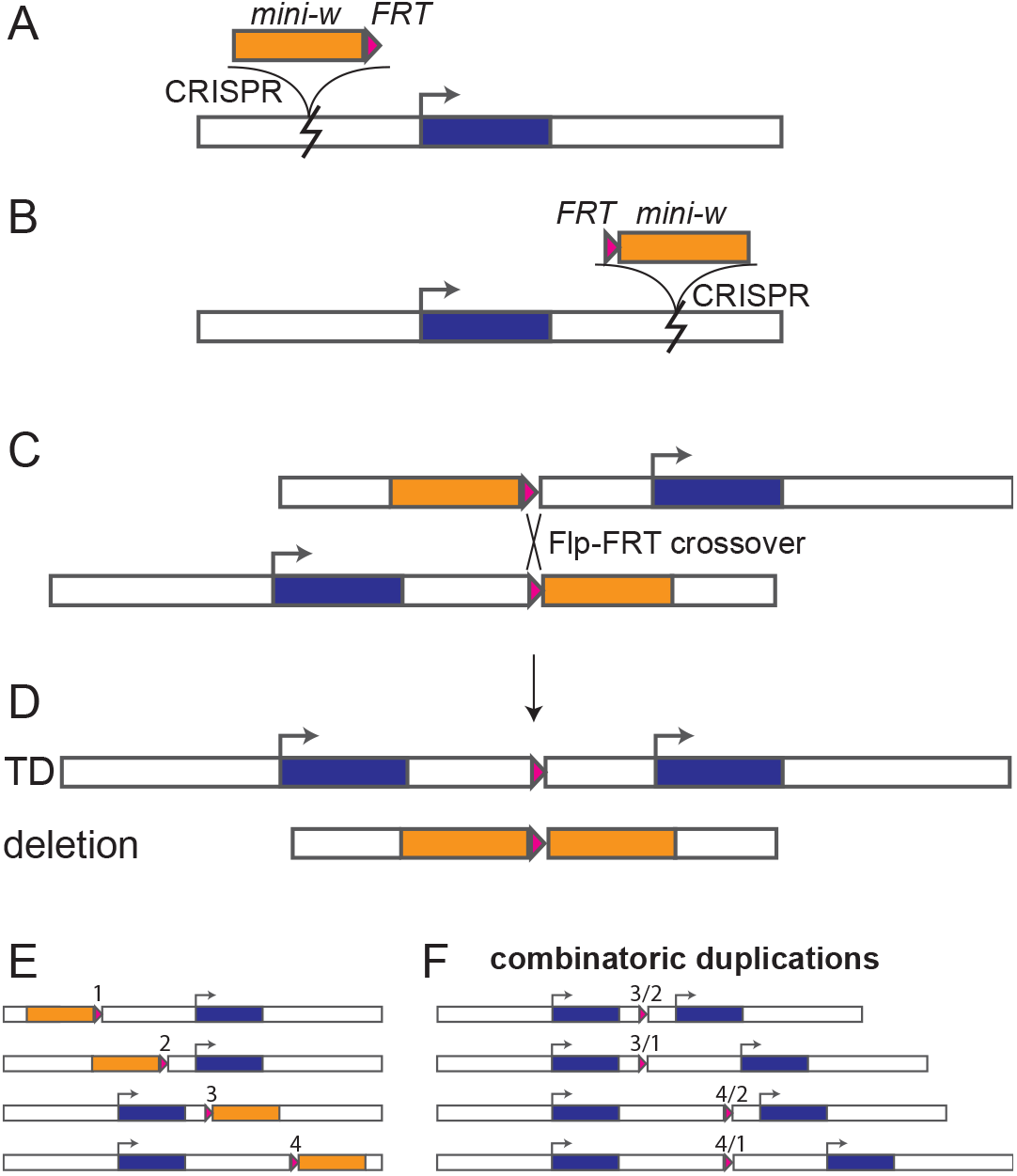
Overview of RMTD procedure. A) marker-FRT and B) FRT-marker constructs are separately inserted on either side of the gene of interest. C) The two marked chromosome homologs are brought together in a heterozygote, along with an unlinked *hs-Flp* recombinase gene. When activated by heat shock, Flp recombinase may cause crossover at the FRT sites. D) Recombinant chromosomes. Note that the tandem duplication (TD) chromosome is unique in lacking a marker gene. E-F) With multiple FRT insertion sites, a variety of duplications of varying structure and size can be engineered.

Serine recombinases, such as Flp, are widely used in genetic engineering to recombine two DNA molecules (Turan and Bode 2011). The Flp enzyme catalyzes a high efficiency of crossover at a specific site, FRT, the Flip Recombination Target. An early Flp-FRT study in *Drosophila* reported the production of various chromosomal rearrangements, including large segmental duplications, using random P-element insertions carrying a FRT site and a *white* (*w^+^*) marker gene (Golic 1994). That study suggested to us that precise tandem duplications of specific genes could be produced if the marker-FRT constructs were targeted to specific sites.

In this paper, we describe the design and production of tandem duplications of the *D. melanogaster Adh* gene (FBgn0000055) using Flp recombinase (Figure 1). Marker-FRT constructs are targeted to specific sites on either side of the gene using CRISPR-Cas9. These constructs are marked with the semi-dominant *mini-w* eye color gene. CRISPR insertions are detected by gain of the *w*^+^ marker. Two such insertions are combined along with a Flp gene, with recombinase-mediated tandem duplications (RMTD) detected by loss of the *w*^+^ markers. We verify that the change in marker phenotype corresponds to the predicted genomic manipulation by quantifying the changes in DNA copy number and ADH enzyme activity. We then discuss practical considerations for design of experiments using this approach.

## Materials and methods

### Fly strains

Genetic manipulations were performed in strains derived from the Cas9-expressing strain BDSC 55821 (Bloomington Drosophila Stock Center #55821, genotype *y[1] M{GFP[E.3xP3]=vas-Cas9.RFP-}ZH-2A w[1118]*). We primarily worked with a culture of this strain, here referred to as BG-55821, obtained from BestGene, Inc. (Chino Hills, CA), in 2018. As described in the Results, this strain was segregating for two distinct *Adh* alleles, of the *Adh^slow^* and *Adh^fast^* types, but this variation was not detected until after most experiments were conducted. For enzyme assays, a culture of BDSC 55821 was obtained in 2022 from the Bloomington Drosophila Stock Center and confirmed to be homozygous for *Adh^slow^*. We also used 55821-Fast, a line derived from BG-55821 that is homozygous for *Adh^fast^*. The source of Flp enzyme was strain BDSC 1929, *y w hs-Flp*; 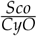. Candidate duplications were isolated using our lab’s balancer stock *y w*; 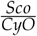.

### CRISPR Insertion sites

Several regions near *Adh* were chosen as insertion sites to place marker-FRT constructs. CRISPR target sites were chosen using the DRSC Find CRISPRs tool https://www.flyrnai.org/crispr/. Sites were chosen if they had an Efficiency Score > 8 and if the sequences of strain BG-55821 matched the reference sequence. Locations of insertion sites MX2, MX5, MX6, and MX10 are shown in Figure 2. Sequence of the candidate insertion site regions from BG-55821 were obtained, as follows. Sequences near the *Adh* gene were obtained using an existing primer set, Adh-clone-F1 and Adh-clone-R1 (Loehlin *et al*. 2019), by PCR amplification, cloning into pGem-T-Easy, and Sanger sequencing with primers listed in Loehlin *et al*. (2019). Sequences of more distal regions, i.e., around sites MX2 and MX10, were obtained Sanger sequence of PCR products using primers listed in File S1.

**Figure 2.**
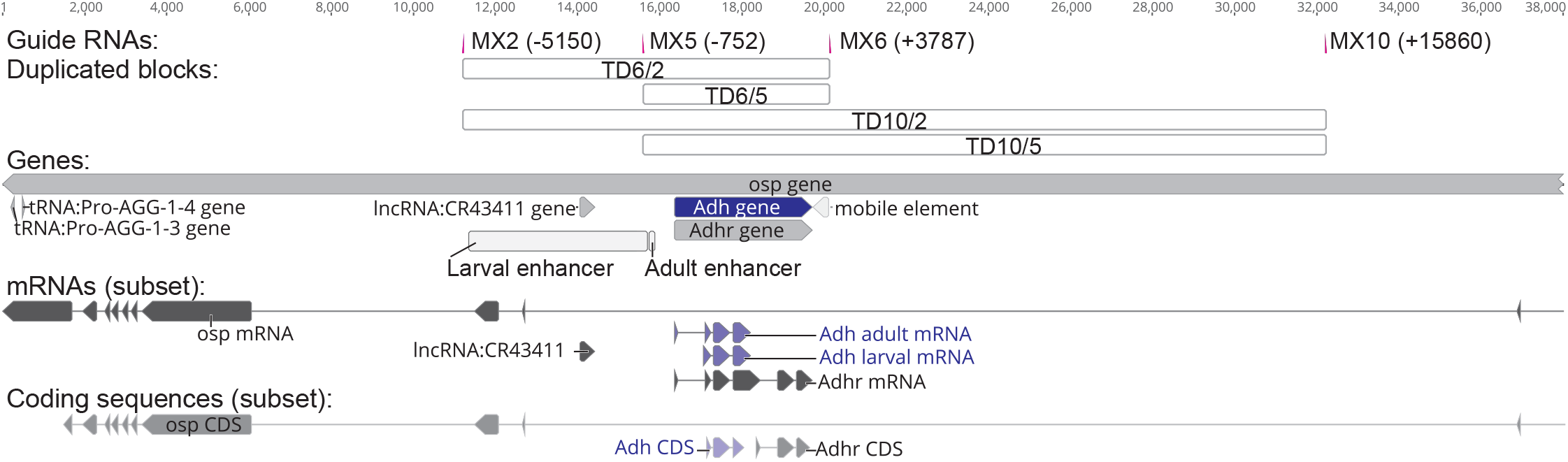
Structure of *Adh* region annotated with CRISPR sites and the predicted span of duplications. Guide RNA sites show the position of insertion for the marker-FRT constructs. Gene features are from the reference annotation, NT_033779. Only a subset of mRNA isoforms is shown for each gene, including the major *Adh* larval and adult isoforms. Enhancer element regions are based on Posakony *et al*. (1985) and Falb and Maniatis (1992). The “mobile element” is part of the reference Iso1 sequence, and is retained for scale, but is not present in any of the strains used here. The element is a 396 bp fragment that replaces a 68bp sequence present in BDSC 55821. The 68bp variant is typical of most other whole genome-sequenced *D. melanogaster* strains (Chakraborty *et al*. 2019).

Guide RNA plasmids were built using the KLD procedure into vector pU6-3-chiRNA (Gratz *et al*. 2014; Dean *et al*. 2022) with primers listed in File S1. Guide sequences are: MX2 CTGAATAATAAGTGGTTGT, MX5 CGAAACCGCTACTCTGGCT, MX6 TAGATGTGCTTAATTATGA, MX10 TTAGCCAGCCAAGATTTAT. G was added at position 20 as in (Gratz *et al*. 2014).

### CRISPaint constructs

Our first approach to CRISPR used the homology-independent CRISPaint approach (Schmid-Burgk *et al*. 2016; Bosch *et al*. 2020). CRISPaint marker-FRT constructs were built using the MoClo (Modular Cloning) approach (Lee *et al*. 2015). Designs were conducted using Geneious Prime (Biomatters, Inc.). Assembly of the w-FRT construct, CRISPaintL, was detailed in (Dean *et al*. 2022). The FRT-w construct, CRISPaintR, was built similarly but using a different Level 1 vector: (t1)ConLS’-(t234r)GFPdropout-(t5)ConRE’-(t67)dmo-miniw- (t8a)AmpRColE1-(t8b)dmo-CRISPaint-targetR. Part sequences are in (Dean *et al*. 2022), except for (t67)dmo-miniw, which has the same sequence as the (t7)miniw but with BsaI overhangs for type 6 (TACA) and type 7 (CCGA), and (t8b)dmo-CRISPaint-targetR, whose sequence is given in File S1. Primers are in File S1.

### HDR-CRISPR assembly

Marker-FRT constructs with homology arms were built according to the plan diagrammed in Figure 3. The design and assembly approaches are detailed in (Dean *et al*. 2022). Briefly, homology-arm PCR products were assembled with a vector fragment and a marker-FRT insert fragment. The same vector, H-arm-CFP, from (Dean *et al*. 2022) was used; this vector is marked with 3xP3-CFP to detect improper insertions. For the marker-FRT inserts, variants of that paper’s FRT-w-FRT insert plasmid were designed. The FRT-w insert plasmid, called Harm-Fw, was built by substituting at type-5 the part (t5)attP39Brc-con2 (primers in File S1). Likewise, the w-FRT insert plasmid, called Harm-wF, was built by substituting the type-1 part (t1)attP39B-con1. Harm-Fw consists of (t1)FRT48-attP39B-(t234)dmo-miniw-(t5)attP39Brc-(t678)KanRColE1. Harm-wF consists of (t1)attP39B-(t234)dmo-miniw-(t5)attP39Brc-FRT48-(t678)KanRColE1.

**Figure 3.**
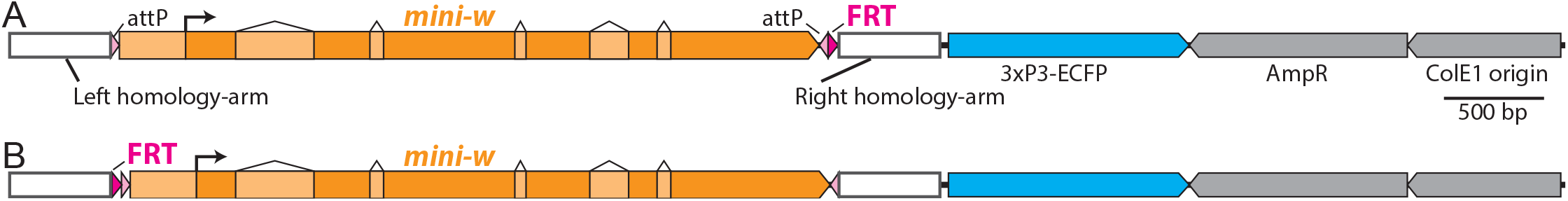
A) Linear structure of an example *w-FRT* construct used for homology-directed-repair, to scale. An additional feature of the construct is the inclusion of PhiC31 attP sites for cassette exchange (Bateman and Wu 2008). B) An example *FRT-w* construct.

The homology arms were PCR amplified from BG-55821 using primers listed in File S1. Because many candidate homology arms contained BsmBI restriction sites, assembly of homology arms to vector fragments was conducted using Gibson assembly, rather than MoClo/Golden Gate assembly. All constructs were verified by Sanger sequencing of junctions and end-to-end coverage of PCR-amplified segments.

### Injections

Plasmids were mixed at a concentration of 500 ng/μL insert and 50 ng/μL each guide RNA. Fly embryo injections were performed by BestGene, Inc. (Chino Hills, CA) into strain BG-55821. Typically, ~300 embryos were injected, then 45-60 G0 flies were crossed to *y w*, then G1 progeny inspected for red eye phenotype. Failed injections were repeated for an additional ~300 embryos. Redeyed progeny were then sib-crossed or balanced using *y w*; 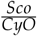 to make homozygous lines. In preparation for tandem duplication, males were crossed to *y w hs-Flp*; 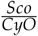 to make homozygous lines that were *y w hs-Flp; *w*^+^ FRT* or *y w hs-Flp; FRT w^+^*.

### Sequence verification of insertions

Correct insertion of marker-FRT constructs was verified by Sanger sequencing of PCR products. For sites MX2 and MX10, insertions were verified using spanning PCR initiated with primers outside the homology arms (primers in File S1). For MX5 and MX6, insertions were verified using junction PCRs from outside the homology arms into *mini-w* (primers in File S1 and Loehlin *et al*. (2019)). To verify that the correct *Adh* allele had been inserted next to, the gene region from each insertion line was PCR amplified using primers Adhseq-1857F/MX6-Rharm-R, then Sanger sequenced. All inserts were found at the correct sites and to be next to the *Adh^slow^* allele of BDSC 55821.

### RMTD crosses

To induce tandem duplication, heterozygous F1 larvae were heat-shocked three times for 1h or 2h in a 37° C microbiological incubator, at days ~2, 4, and 6 after egg laying and then returned to room temperature. As described below, we found the 2h heat shock to be more effective. F1 males were then crossed to balancer *y w*; 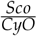 females. To screen for putative tandem duplications, we collected F2 males in separate vials and aged them for 4-9 days to allow the eye color to develop. White-eyed F2 males were then crossed to balancer females to maintain the duplication (“Bal2” refers to either CyO or Sco). The crossing scheme is as follows:

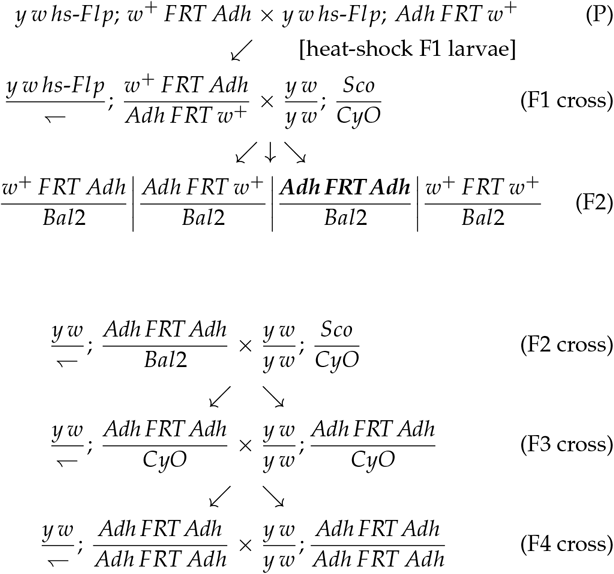

### Copy number determination

Copy numbers of *w* and *Adh* were determined using droplet-digital PCR using a QX200 instrument (Bio-Rad, Inc.). Genomic DNA from single adult male flies was extracted using using the Monarch Genomic DNA Purification Kit (New England Biolabs), using a protocol we developed (dx.doi.org/10.17504/protocols.io.bp2l694qklqe/ v1, Loehlin (2022)). Testing suggested this procedure was more reliable for copy number determination than single-fly “squish” extractions (Gloor and Engels 1992), which are simpler to perform but often showed irregular copy-number calls.

For digital PCR, 2 μL of genomic DNA prep was fragmented by restriction digest in 20 μL reactions with EcoRV-HF and HinDIII-HF (New England Biolabs) for 2h. 4 μL of digest product was assayed in 20 μL PCR reactions using Bio-Rad ddPCR Supermix for Probes (no dUTP) using the manufacturer’s recommended procedure. Assays were duplex, comparing copy number of control gene *RpL32* to *w* or *Adh*. Primers and probes are listed in File S1. ddPCR results were inspected in Bio-Rad QuantaSoft Analysis Pro. Droplets were manually segmented, applying the same threshold to all samples simultaneously. Data were plotted using R package ggplot2.

### Photography

Fly images were recorded on a Zeiss Stemi 305 trinocular stereomicroscope under similar lighting conditions. Flies were killed by freezing for 24h and then photographed within 5 min of thaw to preserve eye color, as in (Dean *et al*. 2022). For the published image, contrast was enhanced by uniformly adjusting levels across images (i.e., one leveling filter was applied to the multipanel figure) using Adobe Photoshop.

### Adh enzyme assay

Adh activity was assayed from 4d old adult male flies, following the high-throughput procedure described in Loehlin *et al*. (2019), using a MultiSkan GO spectrophotometer (Thermo Fisher Scientific). 3 replicate low-density cultures of each genotype were set up and the parents flipped to new vials every 48h. A sample consisted of 4 flies homogenized together; 1-2 samples were measured per vial per day. On a given sampling day (5 day replicates total), all genotypes were measured, though some vials didn’t produce enough flies on certain days for a full set of replicates. ADH enzyme activity (units: Δ*Abs*_340*nm*_ *min*^−1^ *mL*^−1^) and total protein (units: *mg mL*^−1^; Pierce BCA Assay, Thermo Fisher Scientific) of each sample were measured 3 times in technical replicates. Technical replicates were averaged to produce a single response value per homogenate (sample), then log-transformed to account for variance that increased with the mean. The response variable was thus 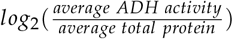. Data were analyzed using a mixed-effects model (R package lme4), with genotype as main effect and vial and day as crossed-factor random effects. Tukey multiple pairwise comparisons were computed from model fits using R package emmeans and presented in the graph using compact letters display (cld) using package multcomp.

## Results

### Development of marker insertion sites for the *Adh* gene

We investigated whether the RMTD approach (Figure 1) was a practical means of generating new tandem duplications from a variety of starting positions. To develop and test the approach, we focused on the model gene *Adh*, whose expression is easily quantified with an enzyme assay. We sought to duplicate a segment containing the *Adh* transcription unit as well as sequences required for expression in adult flies, which have been mapped to within 660bp of the start site of the adult transcript (Posakony *et al*. 1985; Falb and Maniatis 1992). On each side of this segment, we developed CRISPR guide RNAs for two pairs of sites (Figure 2) that could be used to create a range of duplicated blocks. We attempted to insert *w^+^ FRT* constructs on the left of *Adh*, at sites MX2 and MX5, and *FRT w*^+^ constructs on the right, at sites MX6 and MX10. Four possible duplications could be generated from these site combinations, with duplicated block sizes of 4.2 kb, 8.6 kb, 16.3 kb, and 20.7 kb (Figure 2).

### Unsuccessful marker insertion using CRISPaint

Our initial approach to insert marker-FRT sites applied the CRISPaint strategy (Schmid-Burgk *et al*. 2016; Bosch *et al*. 2020; Dean *et al*. 2022). In this approach, the CRISPR/Cas9-induced DNA break in the injected embryo is repaired by nonhomologous end joining. A linearized marker construct is provided, which may insert at the cut site. Marker insertion is detected by phenotypic screening of the offspring of the injected organisms. The marker we used, *mini-w^+^*, is an attenuated version of the gene that partially restores eye pigmentation in *w*^−^ flies within a range from pale yellow to wild-type red that depends on sex, copy number, and the genomic position of insertions (Chetverina *et al*. 2008). In this experiment, no F1 progeny with pigmented eyes were recovered for insertions at sites MX5, MX6, and MX10 (~600 embryos were injected and ~100 G1 families screened per site). The injections targeting site MX2 resulted in two progeny with pigmented eyes. In one line, which had a yellowish eye color in heterozygous males, PCR analysis identified junctions from both the left side and the right side of the genomic DNA into the right side of the marker construct. This suggested that two constructs had inserted in head-to-head orientation. The other line, which had a brownish eye color, did not survive. It remains possible that single insertions of the *mini-w* marker occurred but were not detected due to the weak expression of the marker (described below). Regardless, no correctly oriented insertion lines were identified with this approach.

### Successful marker insertion using HDR-CRISPR

We next attempted to insert marker-constructs into the same sites using the homology-directed-repair (HDR) strategy for CRISPR (Gratz *et al*. 2014), which has worked effectively for us in the past (Loehlin *et al*. 2022; Dean *et al*. 2022). Marker constructs were assembled for each site: either a *w^+^ FRT* or *FRT w^+^* insert flanked with ~500bp homology arms that match the sequence flanking the double-strand-break. The plasmid backbone carried a *3xP3-ECFP* (cyan fluorescent) marker to screen for plasmid backbone insertions, which can be frequent with this procedure (Bier *et al*. 2018; Zirin *et al*. 2021). Per construct, one batch of 300 embryos was injected. One line was recovered at site MX5 and one at site MX10. At site MX6, three independent lines were recovered, with equivalent eye color; one (MX6.1) was chosen for further analysis. At site MX2, three independent lines were recovered. Two lines had darker eyes than the other, suggesting that multiple copies of the marker had inserted, so we chose the lighter-colored line, MX2.3, for further analysis. PCR and sequence analysis of the insertions suggested that each marker-FRT construct had inserted in the correct position and orientation.

Eye color varied among the four insertion lines (Figure 4). Such variation could be the result of 1) multiple insertions of the construct during the homology directed repair process due to crossover repair (Figure 5) (Bier *et al*. 2018) or 2) position-effects from the insertion location. Both factors were evident. Cyan fluorescence was detected in lines MX5.1, MX6.1, and the darker-colored MX2 lines, indicating that the plasmid backbone had inserted. To verify how many copies of the *mini-w*^+^ marker had inserted, we quantified *w^+^* gene copy number using digital PCR analysis (Figure 6). The non-fluorescent MX2.3 and MX10.1 lines contained one inserted *w^+^* copy, as predicted for a canonical HDR-CRISPR insertion. The cyan fluorescent lines MX5.1 and MX6.1 were confirmed to contain two inserted *w^+^* gene copies, as predicted for a backbone insertion at the target site (Figure 5). Position effects of the insertions also appear to play a role in eye color: MX2.3 is more pigmented than MX10.1, though both carry one *w^+^* copy, and MX5.1 is darker than MX6.1, though both carry two *w^+^* copies.

**Figure 4.**
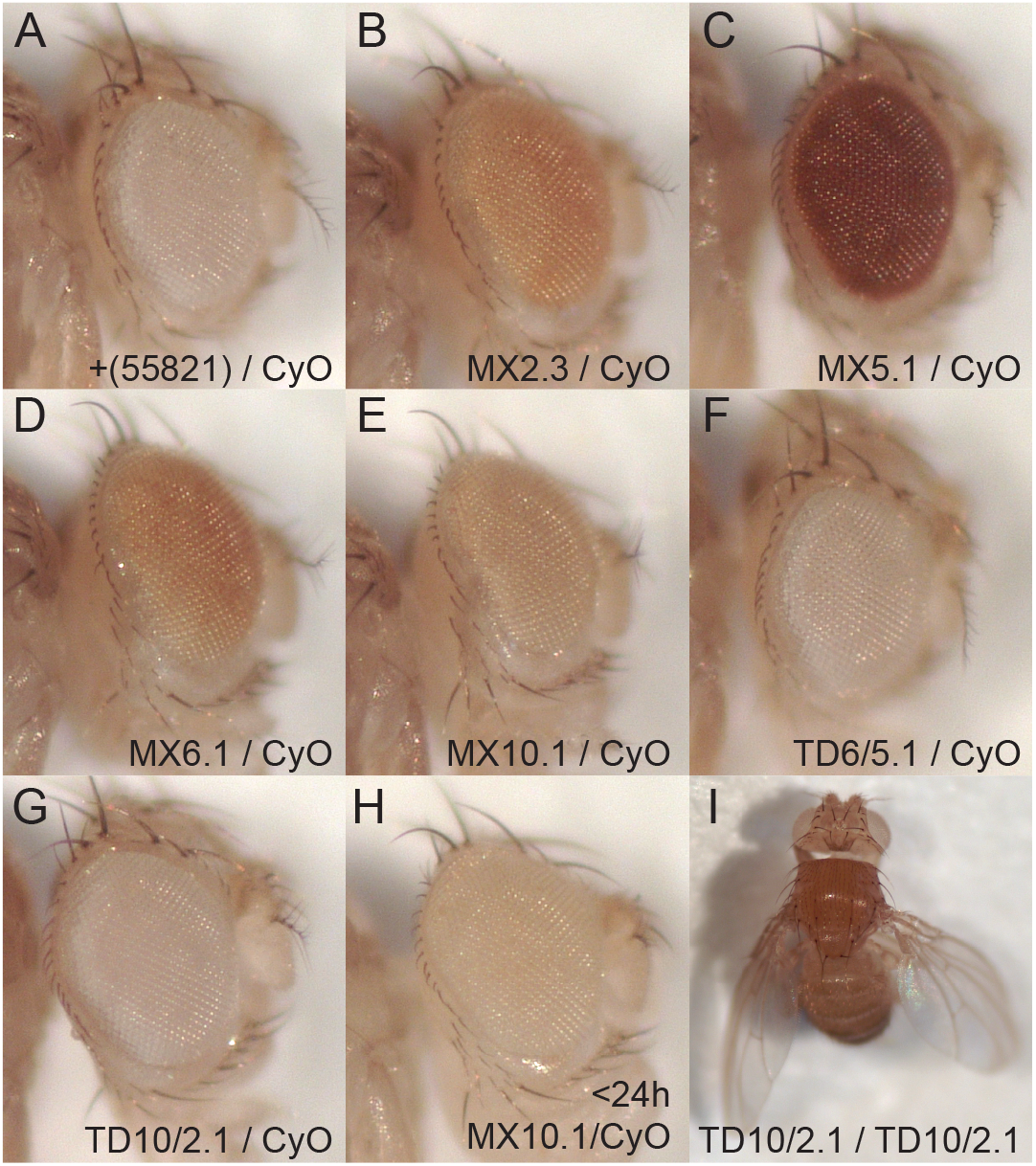
Visible phenotypes used to identify tandem duplications. A-G) Eye color of *w^+^* in the MX insertion lines ranges substantially. Under working conditions, the weak *w^+^* insertions were not easily distinguished from *w^−^*. *w^−^* genotypes include the source stock BG-55821 and tandem duplicates (TD). Hemizygous males aged 6-8d are shown, as they represent the genotype and approximate age when phenotypic screening for tandem duplications was performed. H) Faint eye color of one of the single-copy insertions in a recently eclosed male. I) Outspread/curved wing phenotype of homozygous TD10/2 tandem duplication, also seen in TD6/2, and not in other genotypes.

**Figure 5.**
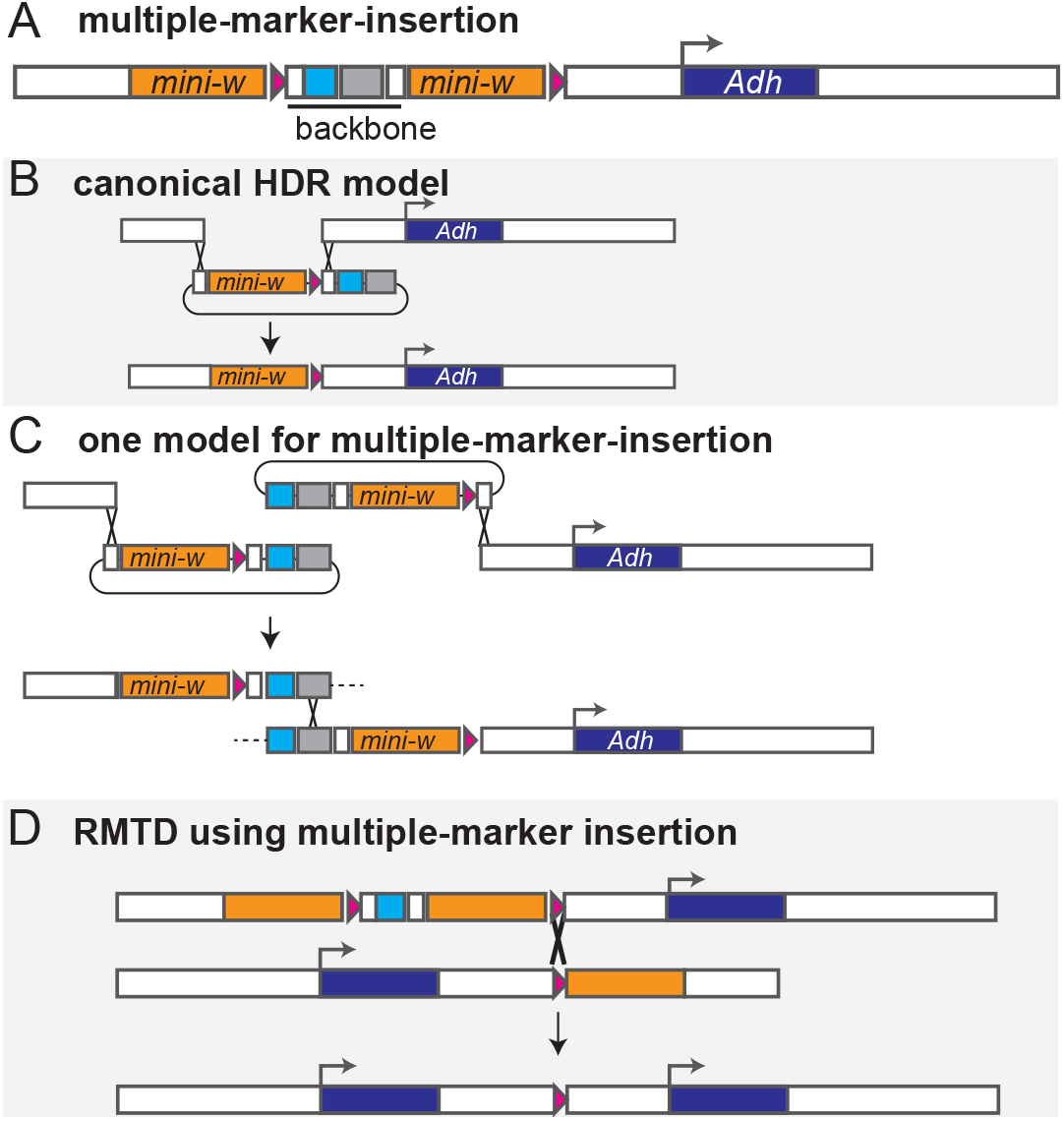
A) Multiple marker insertions are often observed in *Drosophila* HDR-CRISPR experiments. Here, these would contain two *mini-w^+^* and the plasmid backbone, including *3xP3-ECFP* marker (cyan). B) Canonical HDR-CRISPR insertion should insert only the region between the homology arms. C) Alternative model that could explain multiple-marker insertion. Upon double-strand break, each broken chromosome end invades a separate HDR construct. Resolution of an intact chromosome could occur if an additional homologous strand exchange takes place; depending on its position, a multiple-marker-insertion could result. D) Markerless tandem duplication could still be achieved from multiple-marker-insertion(s) if Flp-mediated crossover occurred at the gene-proximal FRT sites.

**Figure 6.**
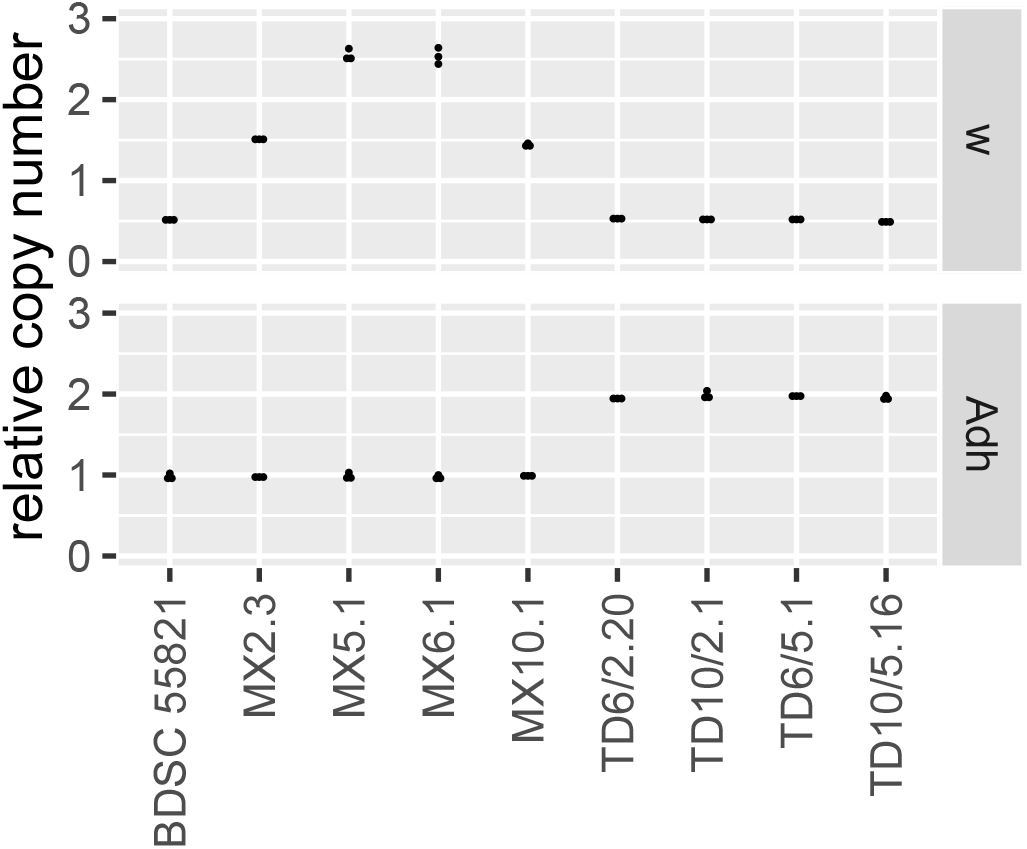
Gene copy number of insertion and tandem duplication lines. Copy number in homozygous male flies was measured in duplex digital PCR assays, with count of focal gene (*w* or *Adh*) normalized to count of autosomal control gene *RpL32*. Each point is a measurement of a separate single-fly genomic DNA preparation, n=3 per genotype. The *w* assay also detects the *w^−^* allele from the endogenous X-linked locus, which is expected to contribute 0.5 copies in these hemizygous males.

Given that the only insertion recovered at two of our sites contained extraneous marker-FRT insertions, it was uncertain whether these would interfere with the RMTD process. We speculated that crossovers at the gene-proximal FRT sites could still recombine out all distal inserted copies, potentially resulting in markerless tandem duplications with the intended structure (Figure 5). If this inference was wrong, we should only be able to obtain markerless tandem duplications from the single insertions (i.e., sites MX2 and MX10).

### Tandem duplications produced

To test the procedure for Flp-mediated duplication, we set up crosses among all four combinations of left-side insertion (*w*^+^ *FRT* at MX2 or MX5) with right-side insertion (*FRT w*^+^ at MX6 or MX10). 8 to 20 F1 males that had been heat-shocked to induce Flp were crossed, singly, to balancer females, then their F2 progeny were screened for loss of eye color. Due to the weak phenotype of single *mini-w* copies, we found that we could only confidently distinguish the diagnostic *w*^−^ phenotype in males, not females, and only after several days of aging (Figure 4).

In our first trial of the duplication procedure, we isolated two independent tandem duplications that combined sites 5 and 6, and chose one line, named TD6/5.1, for further analysis. This confirmed that markerless duplications could be obtained from multiple-marker-insertion lines. We also isolated one duplication line that combined sites 2 and 10, named TD10/2.1. No duplications of 10/5 and 6/2 were recovered in this trial. This rate of tandem duplication recovery was not as high as we had anticipated based on other applications of Flp-FRT in *Drosophila* (e.g. Golic and Lindquist (1989); Harrison and Perrimon (1993); Golic (1994)). Several subsequent trials were fruitless. We speculated that our procedure was suboptimal somehow, and ran across the study of Chou and Perrimon (1992), which determined that germline Flp-FRT activity in *Drosophila* was much higher with longer heat-shock periods, e.g., 2h, versus the 1h heat-shock of typical protocols.

To determine whether a longer heat-shock period would be effective, we repeated the crosses of sites 10 by 5 and 6 by 2, with two heat shocks of 2h applied to F1 larvae. Most F1 adults showed eye color mosaicism, which is an indicator of somatic recombination and thus successful Flp-FRT activity. Approximately 70% of crosses using mosaic-eyed F1 males or females yielded at least one white-eyed F2 male. These observations confirmed that the longer heat-shock was effective. We selected one line of each duplication type, lines TD6/2.20 and TD10/5.16, for detailed analysis.

Outspread wings (Figure 4J) were observed in homozygous TD10/2 and TD6/2 flies, but not other genotypes, consistent with loss of function of *outspread* (*osp*). This makes sense because tandem *Adh* duplications using site MX2 will duplicate two exons of the *osp* gene, resulting in a truncating frame-shift.

To verify that *Adh* had actually been duplicated in the *w^−^* flies, we quantified *Adh* and *w^+^* genomic copy number using digital PCR (Figure 6). Each genotype showed a copy number of *Adh* and *w* genes that was consistent with the RMTD procedure having worked as predicted.

### Tandem duplication increases ADH activity

We are motivated to understand whether tandem duplication will result in a simple doubling of gene expression, or some other outcome, perhaps owing to interactions between the gene copies or other regulatory elements within duplicated blocks. The results presented above suggest that we have produced sequence-identical duplications with different structures, which could begin to address those questions. However, we recovered only a single replicate of several of the genotypes, which limits the explanatory power of a gene expression comparison at present. Nevertheless, we reasoned that we could still conduct a pilot study to determine if the presumed tandem duplications of *Adh* had any effect on gene expression, and if this is uniform or varies in some way with duplication structure, influencing the design of future experiments.

To explore these questions, we compared the expression levels of the marker-insertion and tandem duplication lines described above. We measured expression level using a high-throughput ADH enzyme activity assay that has been tuned to show a one-to-one response to changes in enzyme concentration (Loehlin and Carroll 2016; Loehlin *et al*. 2019). We measured one line each of the four types of duplications and single-copy insertions. To verify that the expression levels of the duplicates were within a normal range, we also measured activity of the pre-insertion starting strain, BDSC 55821 (which carries the *Adh^slow^* allele that was then duplicated) and a single-copy *Adh^fast^* allele, 55821-Fast, in the same genetic background. Typical *Adh^fast^* alleles produce two or more times higher ADH activity than *Adh^slow^* alleles (Laurie *et al*. 1991; Loehlin *et al*. 2019), similar to the anticipated effects of tandem duplicating the *Adh^slow^* allele.

ADH enzyme activity is presented in Figure 7. The marker insertion lines varied in activity: most strikingly, MX5.1 showed 3-fold lower activity than the others. MX2.3 was slightly lower than both MX6.1 and MX10.1 (Tukey’s HSD tests, *P* < 0.05). The two insertions on the right side of *Adh*, MX6.1 and MX10.1, were not significantly different from one another (*P* = 0.98). Compared with the un-inserted strain BDSC 55821, MX2.3 and MX6.1 were similar (*P* = 1.0 and 0.091) but MX10.1 was slightly higher (*P* = 0.011). Variation in the activity of singleton strains might be caused by several factors, such as position effects from the *w*^+^ marker construct, disruption of regulatory elements by the insertion, and variation in genetic background, including from off-target CRISPR mutations.

**Figure 7.**
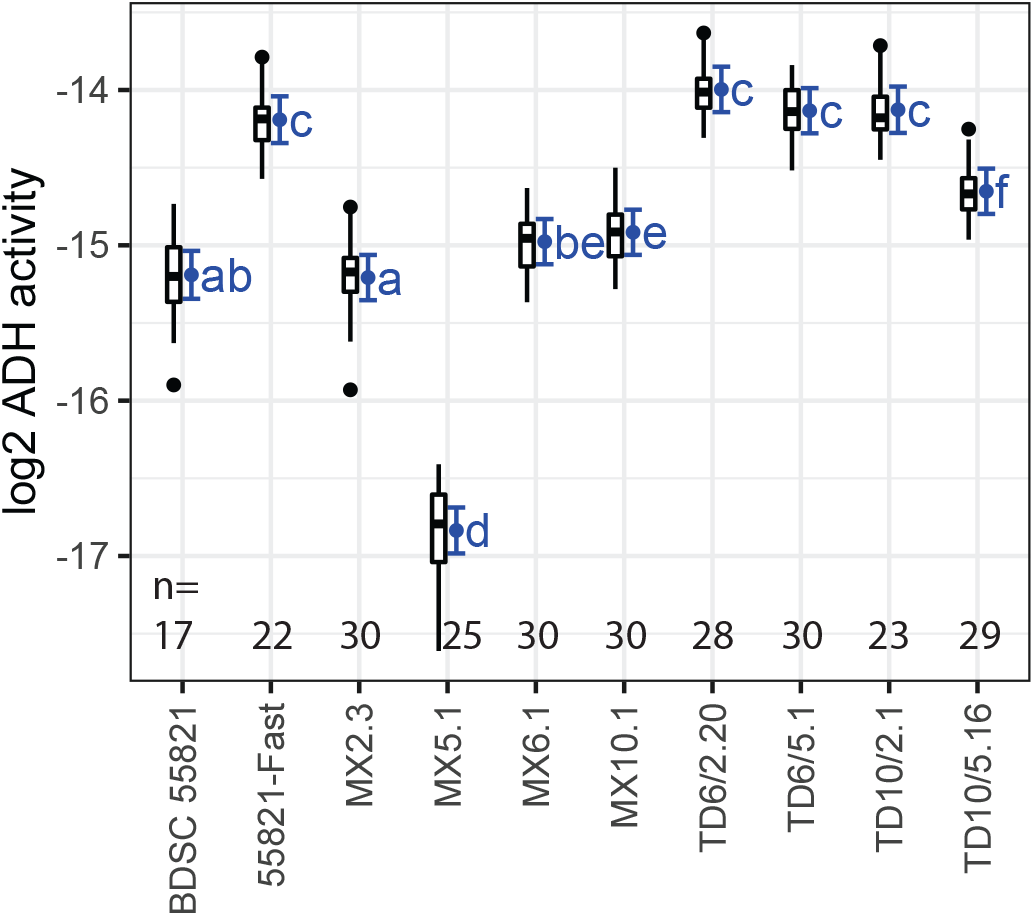
ADH enzyme activity of whole-fly extracts of tandem duplicate and single-copy *Adh* lines. BDSC 55821 has a singlecopy *Adh^slow^* allele that matches the *Adh* sequence in the MX marker-insertion and TD tandem duplicate lines. 55821-Fast has an unrelated *Adh^fast^* allele and is included for comparison. Units are 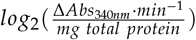, i.e., one unit on the y-axis designates a 2-fold difference in ADH activity. Tukey boxplots (black) show the distribution of data. Blue error bars show mean and 95% confidence intervals from mixed-effects model fit. Tukey HSD multiple comparisons were performed among all pairs of genotypes and are summarized using Compact Letters Display (blue letters). The letters designate groups that are not different, so a genotype labeled ‘ab’ has *P* ≥ 0.05 with any genotype labeled with ‘a’ and/or with ‘b’. *P* < 0.05 if two genotypes do not share any compact letter. n=17-30 replicate extracts were measured per line as indicated, with sample sizes below 30 the result of lower vial productivity.

All four of the tandem duplicates showed significantly elevated expression over the singletons (*P* < 0.05 in each comparison). Three duplicates, TD6/2.20, TD6/5.1, and TD10/2.1, were not significantly different from one another (*P* > 0.4). These were also not different from the *Adh^fast^* strain (*P* > 0.1), supporting our speculation that duplicating an *Adh^slow^* allele would increase activity within the range observed in natural populations. The noteworthy exception among the duplications is TD10/5.16, which showed surprisingly low activity, significantly lower than the other tandems (*P* < 0.0001) but still higher than each *Adh^slow^* singleton (*P* < 0.02). These results demonstrate that the RMTD duplication process increased gene expression of *Adh*. Further, the variation observed raises the possibility that the expression increase might depend on the structure of the duplication. However, we caution that this pilot study is based on limited genotypic replication, so the observed difference among duplicate lines could instead be the result of some other factor such as genetic background.

### Contamination of homology arm sequence

Close analysis of sequences at the end of the project indicated that a contamination had occurred in the creation of the homology constructs. We believe that the impact of this contamination on this experiment was minimal, but we document it here in case this assumption is incorrect and because it has influenced related work in preparation. In a nutshell, we discovered that the culture of BG-55821 used to make the homology arms and the insertion lines carries two segregating *Adh* haplotypes, one *Adh^fast^* and one *Adh^slow^*. The two haplotypes might conceivably have been present at the creation of the strain or might have been introduced subsequently by outcrossing. The two haplotypes present a potential problem for this study because *Adh^fast^* and *Adh^slow^* haplotypes differ substantially in ADH activity (Figure 7). A new culture of BDSC 55821 obtained from the Bloomington stock center in 2022 was found to contain only the *Adh^slow^* haplotype. This culture was used to replace the contaminated BG-55821. Below, we document the effect of the two haplotypes on the insertion lines.

This contamination resulted in a different *Adh* haplotype being used at one stage of the experiment. Specifically, we determined that the aliquot of genomic DNA used to confirm the sequence of guide-RNA sites and to PCR-amplify the homology arms consisted of the *Adh^fast^* haplotype. In contrast, each of the injected HDR constructs had inserted into the *Adh^slow^* haplotype. Once this discrepancy was discovered and understood, PCR and Sanger sequence analysis was used to verify that a single, identical *Adh^slow^* haplotype occurred across the *Adh* transcribed region in BDSC 55821, in each marker-insertion line, and in a slow haplotype that we isolated from BG-55821. This meant that this *Adh^slow^* haplotype could be used as a consistent reference for comparing the single and tandem *Adh* duplications generated here, but with possible contamination of the sequence of the homology arms of the HDR construct.

Using a mismatched sequence in a homology arm could result in incorporation of that mismatch into the chromosome upon repair of the double strand break. Because of the haplotype contamination, each homology arm of the *w-FRT* and *FRT-w* constructs contains a handful of sequence variants from the *Adh^fast^* haplotype. These could have crossed in to the *Adh^slow^* chromosome upon genomic insertion via homologous recombination, or not, depending on where the Holliday junction was situated when the double-strand-break-repair process resolved. To determine whether this had happened, we sequenced the gene-facing homology arm from each marker-insertion line, as any incorporated sequence variants on that side would be retained after tandem duplication. In lines MX2.3, MX5.1, and MX6.1, none of the sequence variants in the gene-facing homology arm crossed in, but in line MX10.1, 4 of 4 fast-type sequence variants in the homology arm had crossed in (Figure S1). It seems unlikely that these variants would influence ADH activity, as the MX10 site is in a position far from *Adh*, and all known expression variation between fast and slow haplotypes has so far been accounted for by variation in the promoter and *Adh* transcribed region (Loehlin *et al*. 2019). In summary, the unanticipated occurrence of segregating *Adh* haplotypes in the BG-55821 strain appears to have a fortuitously minimal impact on the experiments described here.

## Discussion

We successfully created four unique tandem duplications of the *Adh* gene using the RMTD procedure. The resulting duplications increase ADH activity, but to a different degree among lines, raising questions about the structure-expression relationship of tandem duplications. Investigating these questions will require comparison of a broader array of tandem-duplicate structures. Such experiments now appear to be possible. Our experience with developing the RMTD approach suggests several planning considerations for RMTD experiments to be practical and to produce meaningful comparisons.

### Expression of tandem duplicates from varying starting positions

Our pilot enzyme activity study demonstrates that tandem duplications of *Adh* made using RMTD can increase gene expression. Three of four tandem duplications produced about twice the activity of the single-copy lines, while the fourth, TD10/5.16, showed lower expression. The observed variation among tandem duplicate lines is intriguing, in light of the results of past studies that observed deviations from two-fold expression (Loehlin and Carroll 2016; Hayward *et al*. 2017; Rogers *et al*. 2017; Konrad *et al*. 2018; Loehlin *et al*. 2022). Several mechanisms have been hypothesized that could explain why a tandem duplicated gene expresses differently from the sum of two singletons (Loehlin *et al*. 2022). For example, perhaps an enhancer element important for adult expression occurs to the left of site MX5. Insertion of *w-FRT* at MX5 results in lower ADH activity due to separation of the enhancer from the *Adh* promoter. Then, in the duplications from site MX5, the MX5-derived segment would be missing this enhancer, and proper activation of its *Adh* gene would depend on its ability to ‘share’ the enhancer from the left-hand segment. This might explain why the larger duplication TD10/5 has lower activity than the smaller duplication TD6/5. However, the occurrence of an enhancer at this position would be contrary to previous transgene-based mapping (Posakony *et al*. 1985; Corbin and Maniatis 1989), and other explanations are possible.

Overall, we believe that mechanistic interpretation of the expression variation observed here is premature, owing to the limited genotype-level replication available, and given that both single-copy and tandem-duplicate lines varied in activity. The single-copy lines may have varied due to position effects from the *mini-w^+^* marker, and the tandems may have varied due to gene distance or the inclusion of specific regulatory elements, but other explanations are possible. The expression of any particular line is determined by both the experimental manipulation and by unplanned variation among lines, such as off-target CRISPR mutations or variation in the genetic background. Unlinked variation could be partitioned out by increasing replicate creation of specific genotypes, whereas unexpected effects of the experimental manipulation must be controlled by varying the experimental treatment.

From our current perspective, proper assessment of the relationship between single and duplicate expression needs to handle variation arising from both the experimental manipulation and background effects. Discriminating among these factors, in our view, would be best achieved by using a broader variety of insertion sites to generate a range of duplications, and obtaining replicate lines thereof, permitting deviations to be observed multiple times independently and their origins traced. For example, in the enhancer-sharing hypothesis described above, the activity of duplications of *Adh* should show a steep threshold depending on the position of sites near the hypothetical enhancer, but should be nearly invariant among most other combinations of sites. The RMTD approach has the potential to facilitate discovery of such direct effects because it allows combinatoric variation of the position of both ends of a duplication, allowing independent manipulation of both duplicated block size and duplicated block content.

### Practical considerations for design of RMTD experiments

We learned several practical lessons in developing this technique to the present stage. Effective design of the marker-recombinase constructs is critical. In our experience, the weak phenotype of some of the *mini-w* insertions made the phenotypic screening process to be challenging and inefficient, and in retrospect, the weak marker may have also reduced the recovery of CRISPR marker-insertion lines. We were able to resolve the phenotypic ambiguity using a molecular copy-number assay, but a stronger phenotypic marker would have been preferable. One solution might be to use dominant markers such as *y*^+^ or fluorescent proteins. In principle, these would make phenotypic scoring more reliable, and we have conducted preliminary tests that are consistent with this. A stronger version of *mini-w*, perhaps using insulators, might also suffice, and would preserve the additional flexibility gained from semi-dominance of this marker. The version of *mini-w* used here lacks the 3’-flanking *wari* insulator that is present in longer *mini-w* constructs (Chetverina *et al*. 2008), which could have increased the influence of position effects on *w* expression (and, perhaps, *Adh*) that were evident in this study.

The method used to insert the marker-FRT constructs is a central consideration, and this may change as technologies develop. Although we found better success with HDR-CRISPR than CRISPaint for insertion, HDR is more laborious, requiring assembly of a custom construct for each insertion site. The CRISPaint approach remains appealing in that its donor plasmids are universal, requiring only a new guide RNA construct to add a new insertion site. Our insertion success rate with CRISPaint was too low to be useful, which led us to abandon it in favor of the more reliable HDR-CRISPR method. In retrospect, the weak visible marker might have contributed to the low recovery rate, so this method is still worth consideration. The NHEJ insertions produced by this method are less predictable than those made with HDR (Zirin *et al*. 2021), and half of insertions will place the FRT site in a useless reverse-complement orientation. CRISPaint may still be a good idea for a large scale project that targets greater numbers of insertion sites. The HDR-CRISPR approach described here was effective at generating tandem duplications, even with the added obstacles of multiple insertions and a weak phenotypic marker. Kanca *et al*. (2019) recently demonstrated a faster approach to HDR construct assembly using commercially synthesized homology arms that is worth consideration.

Our initial rate of recovery of tandem duplicates was erratic. Increasing the heat shock duration to 2h, as suggested by the experiments of Chou and Perrimon (1992), substantially improved our recovery of replicate tandem duplicates. The sensitivity of this step suggests to us that testing may be needed for other labs to get RMTD to work.

Repeatability and statistical power are major considerations in the design of any experiment. For a genetic-manipulation experiment, obtaining replicate genotypes often poses a practical barrier, especially when increasing effort requires expensive processes such as embryo injection. Here, our ability to draw conclusions about the effects of the duplication on gene expression was limited by only acquiring one replicate of several genotypes. A straightforward solution is to increase the effort to obtain critical genotypes. However, part of the appeal of the RMTD approach is that many different kinds of duplications can be created from a starting set of marker-insertion sites, allowing construction of genotypes that are analogous in structure. For example, this approach could generate similar-sized tandem duplications from a variety of insertion positions, reducing the dependence on any particular genotype for inferring a pattern. The work presented here suggests that such experiments may now be achievable.

Marker removal should be considered in the experimental design if precise quantitative comparisons are required. For the comparison of single and tandem duplicate expression, one needs assurance that no other factors are influencing expression. In this study, we observed varying *Adh* activity among the marker-insertion lines, which might be explained by regulatory interaction (position effects) between *Adh* and *mini-w* or other line-specific effects. If regulatory interactions are the cause, we predict that marker removal should restore a uniform gene expression level among single-copy alleles. One way to achieve marker removal is by recombining a marker-FRT with a FRT-marker construct at the same site. Compared with our current design, this would require a second set of constructs and injections for each site. Alternately, the marker-FRT construct could be redesigned for endogenous marker removal by flanking the marker gene with sites for a different recombinase (e.g., LoxP). This alternate approach would require fewer injections, but would result in a sequence ‘scar’ (LoxP + FRT site) that differs from the scar between tandem duplicates (FRT site alone). It also potentially passages the focal gene through a different genetic background (the Cre recombinase stock), which could introduce a systematic bias between duplicate and non-duplicate lines.

The RMTD procedure can be altered to produce additional genomic manipulations. A straightforward application is targeted deletions of large segments, as these are produced as the counterpart of tandem duplications (Figure 1), replacing the deleted segment with two copies of the marker gene. If markerless deletions are desired, this would require modifying the procedure. One approach to creating a limited set of markerless deletions would be to modify the construct to include a second recombinase site (e.g., LoxP) on the opposite end of the marker from FRT. However, to adapt the RMTD procedure to allow for markerless deletions among any combination of insertion sites (e.g., to delete candidate *cis*-regulatory elements), one needs a pair of marker-insertions where the order of *w*, *FRT*, and gene is reversed relative to the order used for duplication. Thus, obtaining both a marker-FRT and a FRT-marker construct at each insertion site would allow the full range of duplications, deletions, and marker-removals. This strategy would be flexible but imposes a tradeoff, as the resources needed for design, injection, and line maintenance of a second construct at each site might instead be applied to obtaining other insertions, such as expanding the number of insertion sites used for duplication.

## Data availability

Fly strains and plasmids are available from the corresponding author upon request. Sequence of the *Adh* region from strain BDSC 55821 is available at GenBank with the accession numbers: OP794500, OP794501, OP794502.

## Acknowledgments

The authors would like to thank George Yacoub for contributing to an early prototype of this approach and Kyung Shin Kang for designing the *w* and *RpL32* probes for another project. Scott Gratz suggested the 3xP3-CFP backbone-insertion-marker idea to us. Derek Dean provided helpful advice on heat-shocking protocols. Writing Roundtable (Alice Bradley, Catherine Kealhofer, and Katharine Jensen) provided helpful discussions.

## Funding

DWL was supported by startup funds from Williams College and by the National Institute of General Medical Sciences of the National Institutes of Health under Award Number R15GM140429. MX was supported by a Summer Science Research fellowship from Williams College. GLM, RK, and ER were supported through summer research fellowships under the NIH award. The content is solely the responsibility of the authors and does not necessarily represent the official views of the National Institutes of Health.

## Conflicts of interest

The authors declare no conflicts of interest.

## Notes

### Competing Interest Statement

The authors have declared no competing interest.

## Literature cited

Ashburner M, Misra S, Roote J, Lewis S, Blazej R, Davis T, Doyle C, Galle R, George R, Harris N et al. 1999. An exploration of the sequence of a 2.9-Mb region of the genome of *Drosophila melanogaster*: the *Adh* region. Genetics. 153:179–219.

Bateman JR, Wu CT. 2008. A simple polymerase chain reaction-based method for the construction of recombinase-mediated cassette exchange donor vectors. Genetics. 180:1763–1766.

Bier E, Harrison MM, O’Connor-Giles KM, Wildonger J. 2018. Advances in engineering the fly genome with the CRISPR-Cas system. Genetics. 208:1–18.

Birchler JA, Yang H. 2022. The multiple fates of gene duplications: deletion, hypofunctionalization, subfunctionalization, neofunctionalization, dosage balance constraints, and neutral variation. The Plant Cell..

Bosch JA, Colbeth R, Zirin J, Perrimon N. 2020. Gene knock-ins in *Drosophila* using homology-independent insertion of universal donor plasmids. Genetics. 214:75–89.

Cardoso-Moreira M, Arguello JR, Gottipati S, Harshman LG, Grenier JK, Clark AG. 2016. Evidence for the fixation of gene duplications by positive selection in *Drosophila*. Genome Research. 26:787–798.

Carvalho CM, Lupski JR. 2016. Mechanisms underlying structural variant formation in genomic disorders. Nature Reviews Genetics. 17:224–238.

Chakraborty M, Emerson J, Macdonald SJ, Long AD. 2019. Structural variants exhibit widespread allelic heterogeneity and shape variation in complex traits. Nature Communications. 10:1–11.

Chetverina D, Savitskaya E, Maksimenko O, Melnikova L, Zaytseva O, Parshikov A, Galkin AV, Georgiev P. 2008. Red flag on the *white* reporter: a versatile insulator abuts the *white* gene in *Drosophila* and is omnipresent in *mini-white* constructs. Nucleic acids research. 36:929–937.

Chou TB, Perrimon N. 1992. Use of a yeast site-specific recombinase to produce female germline chimeras in *Drosophila*. Genetics. 131:643–653.

Corbin V, Maniatis T. 1989. The role of specific enhancer-promoter interactions in the drosophila adh promoter switch. Genes & development. 3:2191–2200.

Dean DM, Deitcher DL, Paster CO, Xu M, Loehlin DW. 2022. “a fly appeared”: *sable*, a classic *Drosophila* mutation, maps to *Yippee*, a gene affecting body color, wings, and bristles. G3. 12:jkac058.

Falb D, Maniatis T. 1992. A conserved regulatory unit implicated in tissue-specific gene expression in *Drosophila* and man. Genes & development. 6:454–465.

Gloor G, Engels W. 1992. Single fly preps for PCR. Drosophila Information Service. 71:148–149.

Golic KG. 1994. Local transposition of P elements in *Drosophila melanogaster* and recombination between duplicated elements using a site-specific recombinase. Genetics. 137:551–563.

Golic KG, Lindquist S. 1989. The FLP recombinase of yeast catalyzes site-specific recombination in the *Drosophila genome*. Cell. 59:499–509.

Gratz SJ, Ukken FP, Rubinstein CD, Thiede G, Donohue LK, Cummings AM, O’Connor-Giles KM. 2014. Highly specific and efficient CRISPR/Cas9-catalyzed homology-directed repair in *Drosophila*. Genetics. 196:961–971.

Harrison DA, Perrimon N. 1993. Simple and efficient generation of marked clones in *Drosophila*. Current Biology. 3:424–433.

Hayward CP, Liang M, Tasneem S, Soomro A, Waye JS, Paterson AD, Rivard GE, Wilson MD. 2017. The duplication mutation of Quebec platelet disorder dysregulates PLAU, but not C10orf55, selectively increasing production of normal PLAU transcripts by megakaryocytes but not granulocytes. PLoS One. 12:e0173991.

Kanca O, Zirin J, Garcia-Marques J, Knight SM, Yang-Zhou D, Amador G, Chung H, Zuo Z, Ma L, He Y et al. 2019. An efficient CRISPR-based strategy to insert small and large fragments of DNA using short homology arms. Elife. 8:e51539.

Konrad A, Flibotte S, Taylor J, Waterston RH, Moerman DG, Bergthorsson U, Katju V. 2018. Mutational and transcriptional landscape of spontaneous gene duplications and deletions in *Caenorhabditis elegans*. Proceedings of the National Academy of Sciences. 115:7386–7391.

Lan X, Pritchard JK. 2016. Coregulation of tandem duplicate genes slows evolution of subfunctionalization in mammals. Science. 352:1009–1013.

Laurie CC, Bridgham JT, Choudhary M. 1991. Associations between DNA sequence variation and variation in expression of the *Adh* gene in natural populations of *Drosophila melanogaster*. Genetics. 129:489–499.

Lee ME, DeLoache WC, Cervantes B, Dueber JE. 2015. A highly characterized yeast toolkit for modular, multipart assembly. ACS Synthetic Biology. 4:975–986.

Levo M, Raimundo J, Bing XY, Sisco Z, Batut PJ, Ryabichko S, Gregor T, Levine MS. 2022. Transcriptional coupling of distant regulatory genes in living embryos. Nature. 605:754–760.

Loehlin DW. 2022. Drosophila genomic DNA isolation using NEB Monarch kit. https://dx.doi.org/10.17504/protocols.io.bp2l694qklqe/v1.

Loehlin DW, Ames JR, Vaccaro K, Carroll SB. 2019. A major role for noncoding regulatory mutations in the evolution of enzyme activity. Proceedings of the National Academy of Sciences. 116:12383–12389.

Loehlin DW, Carroll SB. 2016. Expression of tandem gene duplicates is often greater than twofold. Proceedings of the National Academy of Sciences. 113:5988–5992.

Loehlin DW, Kim JY, Paster CO. 2022. A tandem duplication in *Drosophila melanogaster* shows enhanced expression beyond the gene copy number. Genetics. 220:iyab231.

Posakony J, Fischer J, Maniatis T. 1985. Identification of dna sequences required for the regulation of drosophila alcohol dehydrogenase gene expression. In: .volume 50. pp. 515–520. Cold Spring Harbor Laboratory Press.

Rogers RL, Shao L, Thornton KR. 2017. Tandem duplications lead to novel expression patterns through exon shuffling in *Drosophila yakuba*. PLoS Genetics. 13:e1006795.

Schmid-Burgk JL, Höning K, Ebert TS, Hornung V. 2016. CRISPaint allows modular base-specific gene tagging using a ligase-4-dependent mechanism. Nature communications. 7:1–12.

Turan S, Bode J. 2011. Site-specific recombinases: from tag-and-target-to tag-and-exchange-based genomic modifications. The FASEB Journal. 25:4088–4107.

Zirin J, Bosch J, Viswanatha R, Mohr SE, Perrimon N. 2021. State-of-the-art CRISPR for in vivo and cell-based studies in *Drosophila*. Trends in Genetics..

