## Supplementary material for "Demonstration of *in vivo* engineered tandem duplications of varying sizes using CRISPR and recombinases in *Drosophila melanogaster*": Figure S1

```
NT_033779          AGAGGTGTTGCACTTTTGTATTTATTTATGAATGAAACTGACAGAGACAAAGAGACGGCC   60
BDSC 55821         ............................................................   60
BG55821 (fast)     .......N........................N...............NNN...N..A..   60
HarMX10 construct  .........................................................A..   60
MX10.1 PCR product .........................................................A..   60

NT_033779          TGGTCGTGGAGGTGGAGAAGGAGGCTGGCAGGGAGTGGGAAAGTTTCAAGTGCACTGAGA  120
BDSC 55821         ............................................................  120
BG55821 (fast)     .............N......N.......................................  120
HarMX10 construct  ............................................................  120
MX10.1 PCR product ............................................................  120

NT_033779          AAAATTTTAAGTAAATTAAAAGCAATATTTTGCGTATACAAATTGCTCATACATATAATC  180
BDSC 55821         ............................................................  180
BG55821 (fast)     ............................................................  180
HarMX10 construct  ............................................................  180
MX10.1 PCR product ............................................................  180

NT_033779          CTCAAAAAATGAATTGTTTTCGTTAACTCAAAAGAATGTTACAATATGTACTGCCTAACT  240
BDSC 55821         ............................................................  240
BG55821 (fast)     ......................................................T.....  240
HarMX10 construct  ......................................................T.....  240
MX10.1 PCR product ......................................................T.....  240

NT_033779          TATGTTTGGCATTTTGCATTTTTTGTCAACTTTTTCTGTCAGTGTGCAAATCTGCAATAT  300
BDSC 55821         ............................................................  300
BG55821 (fast)     ..............-....................................N........  299
HarMX10 construct  ..............-.............................................  299
MX10.1 PCR product ..............-.............................................  299

NT_033779          GCATCTCGTACTCATATGCAAGCCAAGGTCGATTTCTGTTCCGTTCAACTGCAGCAGCAG  360
BDSC 55821         ............................................................  360
BG55821 (fast)     ....................A.......................................  419
HarMX10 construct  ....................A.......................................  419
MX10.1 PCR product ....................A.......................................  419

NT_033779          TTtattgg                                                      422
BDSC 55821         ..tattgg                                                      422
BG55821 (fast)     ..tattgg                                                      421
HarMX10 construct  ..        [FRT w+]                                            421
MX10.1 PCR product ..                                                            421
```

**Figure S1.** Alignment of MX10 left homology arm sequence, showing that the homology arm was derived from *Adh*$^{fast}$ sequence, and that these sequence variants incorporated into the chromosome in the MX10.1 insertion line. Sanger sequences were generated from plasmid of construct HarmX10, from PCR product with primers 12500-1F/12500-1R (DNA from BDSC 55821 and BG55821), or PCR product with primers 12500-1F/uwseq-R6 on DNA from MX10.1. 'N' refers to an ambiguous base call – the BG55821 sequence read was low quality but clearly matched MX10 at the variant bases. The MX10 guide RNA site is marked with bold.
